# Transcriptome analyses reveal tau isoform-driven changes in transposable element and gene expression

**DOI:** 10.1101/2021.04.30.442101

**Authors:** Jennifer Grundman, Brian Spencer, Floyd Sarsoza, Robert A. Rissman

## Abstract

Alternative splicing of the gene MAPT produces several isoforms of tau protein. Overexpression of these isoforms is characteristic of tauopathies, which are currently untreatable neurodegenerative diseases. Though non-canonical functions of tau have drawn interest, the role of tau isoforms in these diseases has not been fully examined and may reveal new details of tau-driven pathology. In particular, tau has been shown to promote activation of transposable elements — highly regulated nucleotide sequences that replicate throughout the genome and can promote immunologic responses and cellular stress. This study examined tau isoforms’ roles in promoting cell damage and dysregulation of genes and transposable elements at a family-specific and locus-specific level. We performed immunofluorescence, Western blot and cytotoxicity assays, along with paired-end RNA sequencing on differentiated SH-SY5Y cells infected with lentiviral constructs of tau isoforms and treated with amyloid-beta oligomers. Our transcriptomic findings were validated using publicly available RNA-sequencing data from Alzheimer’s disease, progressive supranuclear palsy and control human samples from the Accelerating Medicine’s Partnership for AD (AMP-AD).

Significance for biochemical assays was determined using Wilcoxon ranked-sum tests and false discovery rate. Transcriptome analysis was conducted through DESeq2 and the TEToolkit suite available from the Hammell lab at Cold Spring Harbor Laboratory. Our analyses show overexpression of different tau isoforms and their interactions with amyloid-beta in SH-SY5Y cells result in isoform-specific changes in the transcriptome, with locus-specific transposable element dysregulation patterns paralleling those seen in patients with Alzheimer’s disease and progressive supranuclear palsy. Locus-level transposable element expression showed increased dysregulation of L1 and Alu sites, which have been shown to drive pathology in other neurological diseases. We also demonstrated differences in rates of cell death in SH-SY5Y cells depending on tau isoform overexpression. These results demonstrate the importance of examining tau isoforms’ role in neurodegeneration and of further examining transposable element dysregulation in tauopathies.

## Introduction

Tauopathies are a class of neurodegenerative diseases for which there are currently no symptomatic or curative treatments. Although tauopathies are broadly characterized by accumulation of pathological tau protein, the extent and isoform expression of tau varies across diseases [15]. Tau protein exists in six isoforms in humans, formed from the alternative splicing of the gene MAPT, located on chromosome 17. Tau isoforms are distinguished by number of N-terminal repeats (0, 1, or 2) and microtubule-binding repeats (3 or 4; these are respectively referred to as 3R or 4R tau isoforms) [9]. Tau isoform imbalance is thought to be mechanistically linked to neurodegenerative diseases, such as Alzheimer’s disease (AD) (3R and 4R tau), progressive supranuclear palsy (PSP) (4R tau), and Pick’s disease (PiD) (3R Tau) [15], with recent studies demonstrating that imbalance in tau isoform expression can lead to different forms of neurodegeneration [1, 14, 27].

While most tau research focuses on tau’s stabilization of microtubules along with its aggregation into intracellular neurofibrillary tangles (NFTs), tau has significant roles in other cell functions that are affected in neurodegeneration [9, 29]. Recent studies show that tau protein may protect DNA and RNA from heat stress and dysfunction of tau may be linked to heterochromatin relaxation, transcriptomic changes, and disruption of normal protein synthesis patterns [7, 9, 12, 19, 32, 37].

Tau-influenced epigenetic and transcriptomic changes may also account for increased transposable element (TE) expression observed in some tauopathies. Two recent studies found tau expression correlating with TE expression in diseases such as AD and PSP [8, 33]. TEs are nucleotide sequences that can relocate throughout the genome either by copying themselves through an RNA intermediate and inserting these copies into other genomic regions (i.e., “copy and paste”) or by simply translocating to a new genomic area (i.e., “cut and paste”) [24]. Thus, TEs may cause insertion mutations and carry out regulatory functions, and may promote inflammation through creating increased interferon responses [13, 20, 34]. Notably, treatment of tau-transgenic Drosophila with reverse transcriptase inhibitors to suppress TEs can result in marked phenotypic improvement and reduced cell death, presenting a potentially exciting new avenue for therapeutic approaches [33].

To determine whether tau isoforms differ in the way they influence cell dysfunction, we differentiated and infected neuroblastoma SH-SY5Y cells with lentiviruses expressing 3R or 4R or a combination of both tau isoforms. We then tested whether tau isoform overexpression induced changes in tau localization, DNA damage, cell death, and the transcriptome, including changes in TE expression. Because AD is the most prevalent neurodegenerative disease, we also characterized how these changes could be modulated by amyloid beta (Aβ). To validate our *in vitro* model, we repeated our RNA-sequencing (RNA-seq) analysis on publicly available human data from AD, PSP, and non-demented patients. Our study provides new data suggesting a tau isoform-dependent difference in TE activation and repression and identifies the genomic locations of activated and repressed TEs.

## Materials and Methods

### Construction of lentivirus vectors

The human 3RTau (0N3R, 352) [L266V, G272V] cDNA [25] or 4RTau (1N4R, 412) cDNA Open Biosystems were PCR amplified and cloned into the third-generation self-inactivating lentivirus vector [36] with the CMV promoter driving expression producing the vector. Lentiviruses expressing tau (LV-3Rtau, LV-4Rtau), and empty vector (LV-Ctrl) were prepared by transient transfection in 293T cells [36]. For further details on LV-Ctrl production, see previously described methods [30].

### Cell culture

For these experiments, we used the SH-SY5Y human neuroblastoma cell line, which expresses normal levels of human tau and has been used as an *in vitro* system for modeling neurodegenerative diseases [38]. Undifferentiated SY5Y cells underwent fewer than 20 passages, were passaged weekly, and were maintained with media composed of 10% fetal bovine serum and a 1:1 mixture of Eagle’s Minimum Essential Medium (MEM) and Ham’s F12 (F12). In all experiments, cells were infected at plating at a multiplicity of infection (MOI) of 20, and media was replaced the morning after plating. Cells were infected with LV-Ctrl, LV-3Rtau, LV-4Rtau, or an equal ratio of LV-3Rtau and LV-4Rtau.

Cells were differentiated for 9 days with media composed of 15 nM Retinoic Acid (Sigma), 3% fetal bovine serum, and 1:1 MEM:F12, which was replaced every 1–2 days. On the 8th day, 24 hours before collection, all cells were treated with either 50 nM oligomeric Aβ (Aβ42, rPeptide, cat no: A-1163), prepared according to previously published methods [31] or vehicle (DMSO). For a schema of the cell culture infection and treatment procedure, see Fig 1B.

**Fig 1.**
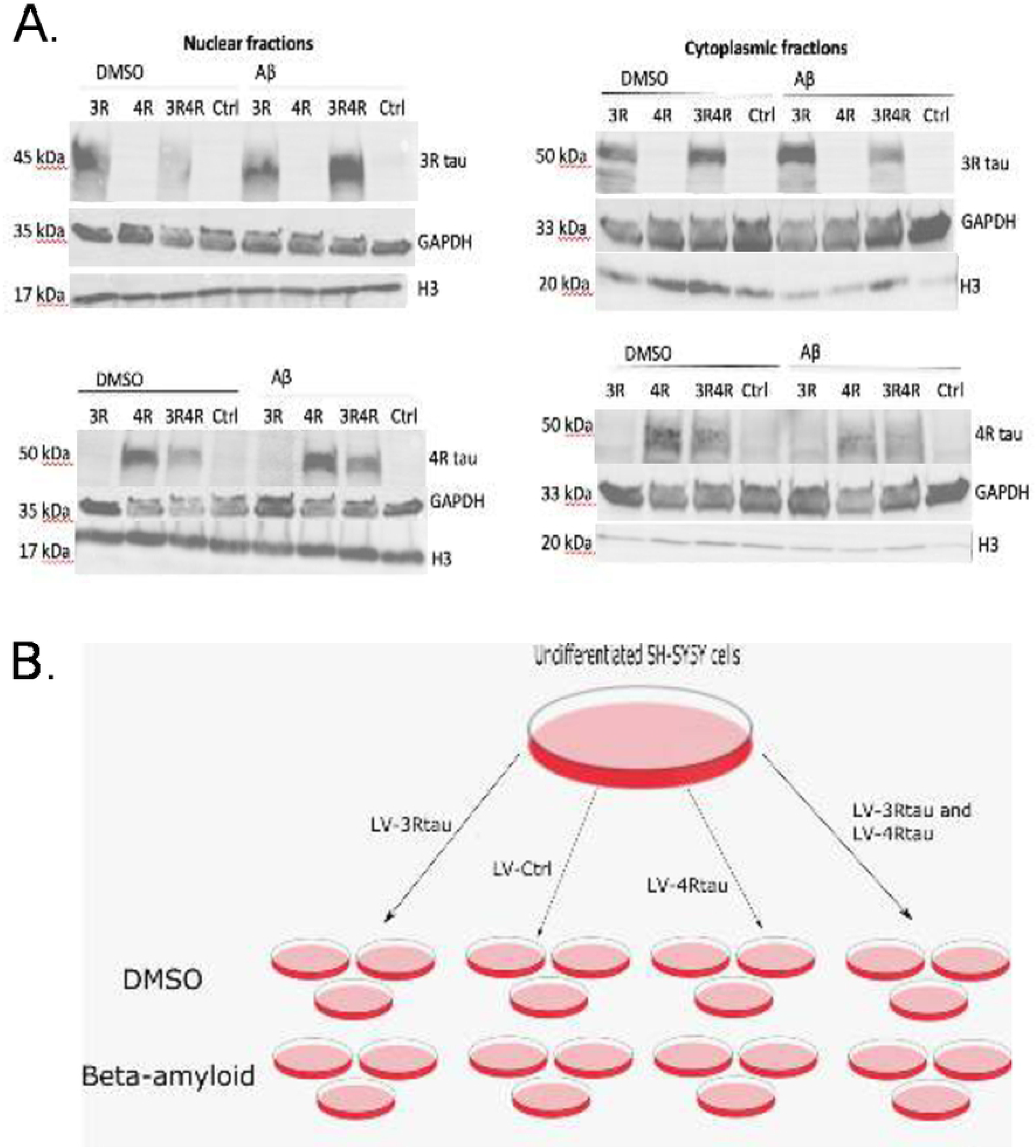
Validation of a typical infection and treatment workflow. (A) Use of tau isoform-specific antibodies in Western blotting demonstrates tau isoform-specific overexpression in both nuclear and cytoplasmic fractions as a result of infection with tau lentiviruses (N=1). (B) A typical workflow for infecting cells with N=3. Cells were treated with 50nM Aβ or DMSO 24 hours before collection.

### Subcellular fractionation and Western blot

Cells were plated onto 10 cm dishes and were treated as previously described. After 24 hours of incubating with Aβ, cells were collected and separated into nuclear and cytoplasmic fractions according to manufacturer recommended subcellular fractionation protocol (https://www.abcam.com/protocols/subcellular-fractionation-protocol). After protein concentration for all fractions were determined using a reducing agent-compatible BCA assay (ThermoFisher, cat no. 23252), 12.1 ug of protein from each sample was loaded onto a 10% Tris-Glycine gel (Biorad, cat no. 4561034) and then transferred onto a PVDF membrane and blocked for one hour with 5% BSA in 0.1% Tween/1xTBS. Membranes were probed with primary antibodies 4R tau (Millipore Sigma, cat no. 05-804), 3R tau (Millipore Sigma, cat no. 05-803), GAPDH (Abcam, cat no. 181602), and Histone 3 (Abcam, cat no. ab1791) overnight followed by the respective secondary HRP antibodies (mouse or rabbit) at 1:5000 and imaged using West Pico Supersignal (ThermoFisher, cat no. 34580)

### Immunofluorescence and confocal microscopy

Cells were plated onto glass poly L-lysine-coated coverslips into 12-well cell culture plates and infected and differentiated as described previously. Following Aβ or vehicle treatment, coverslips were washed once with ice-cold 1xPBS and fixed with ice-cold 4% PFA.

All combinations of LV-tau and Aβ/DMSO treatment were assessed via immunofluorescent staining and confocal microscopy. Coverslips were washed with 1xPBS, followed by 1 hour of blocking with 3% normal goat serum in 0.2% Triton-X/1xPBS. Samples were then incubated with a primary antibody followed by appropriate secondary anti-rabbit Alexa Fluor 568 secondary antibody before being counterstained with DAPI and mounted. The following primary antibodies were used: 1:500 anti-rabbit yH2AX (Bethyl Laboratories, cat no. A300-081A-M) and 1:500 polyclonal rabbit, anti-human tau (Dako, cat no. A0024).

All slides were imaged with a DMI 4000B inverted fluorescent microscope (Leica, Germany) with an attached TCS SPE confocal system (Leica), using a Leica 63X (N.A. 1.3) objective. Analysis of images was carried out using Fiji. For samples stained for yH2AX, nuclei were selected using the DAPI/blue channel and traced by hand, excluding nuclei cut off by the edges of the image or that overlapped with each other or with neurites; nuclear foci were quantified using an online protocol provided by Duke University Microscopy Core with a maxima threshold determined using secondary-only control images in order to distinguish true signal from background. For samples stained for tau, cells were segmented by hand into nuclear (with DAPI staining as a reference) and cytoplasmic (defined as the area of the cell outside the nucleus) regions and corrected total cell fluorescence (CTCF) was calculated each region. The ratio of CTCF coming from the nucleus versus the cytoplasm was then assessed to determine total tau localization within the cell. Statistical significance for both datasets was assessed using a Wilcoxon ranked-sum test and multiple testing corrections were done using the False Discovery Rate. Figures were created with the R package ggplot2.

### Cytotoxicity

The CellTox™ Green Cytotoxicity Assay (Promega, cat no. G8742) was used according to the manufacturer’s instructions to assess cytotoxicity as the result of LV-tau infection and Aβ treatment. Cells were plated and infected into three 96-well cell culture plates. Because the assay allows for fluorescence to be measured for up to 72 hours, each plate was used to measure cell death over a period of 2–3 days, for a total of 9 days (the time period of differentiation for all cells). The first plate was designed to measure cell death resulting from infection over the first 3 days after plating (the assay was started after media was replaced following plating) (N=14), the second plate for days 4–6 after plating and infection (N=14), and the third plate was used to measure cell death as a result of infection and Aβ treatment over days 8–9 (N=7). Cells were fed every 72 hours or when the assay for each plate was performed.

Figures were generated using the R package ggplot2, and statistical significance was assessed using a Wilcoxon ranked-sum test, with multiple testing corrections done using the False Discovery Rate.

### RNA isolation and library preparation

Cells were plated and infected in triplicate into 6-well cell culture plates. Following treatment with Aβ-42 or vehicle (DMSO), cells were scraped from the plates and their RNA was collected using TRIzol (Invitrogen, cat no. 10296010) according to the manufacturer’s instructions.

RNA quality checking, library preparation and sequencing were conducted at the IGM Genomics Center, University of California, San Diego, La Jolla, CA. All RNA had at least an RNA integrity score of 7.9, with the majority of samples scoring greater than 9.5. Libraries were constructed using poly(A) selection to generate 100 bp paired-end reads and were sequenced with an Illumina NovaSeq 6000 that was purchased with funding from a National Institutes of Health SIG grant (#S10 OD026929).

Publicly available RNA-sequencing data from AD, PSP, and control patients (obtainable through AMP-AD Knowledge Portal, doi:10.1038/sdata.2016.89) were also analyzed. For details on how this data was generated, see Allen et al., 2016 [2].

### RNA-seq processing and analysis

Sequencing quality for all FASTQ files was obtained via FastQC [3] both before and after adapter removal using BBduk (BBMap_38.73). FASTQ files were then mapped to the GRCh38 human reference genome and GTF (release number 98) available through Ensembl using STAR (version 2.7.3a) [4]. Qualimap (v2.2.1) [22] was used to visualize the quality of resulting bam files. Gene counts were determined through featureCounts from the package Subread (version 1.6.4) [16], differential gene expression was computed using DESeq2 (version 1.26.0) [17], and GO analysis was carried out using clusterProfileR [39].

TEcount (2.0.3) and TElocal (0.1.0), from the Hammell lab’s TEToolkit suite, were used to determine transposable element expression. TEcount [11] was used in order to produce a global view of TE family expression and TElocal to show localized TE expression, following the alignment protocol and settings described by the authors [10], with the maximum number of iterations set to 100 (the default).

Publicly available FASTQ data generated from the temporal cortex of AD, PSP, PA, and non-demented control patients obtainable through the AMP-AD Knowledge Portal (doi:10.1038/sdata.2016.89) were downloaded from Synapse using the R package synapser (https://github.com/Sage-Bionetworks/synapser). These FASTQ files were selected based on RNA integrity number (>8.0) and quality assessment with FastQC to yield a total analyzed sample size of 60 AD, 71 PSP, and 33 control patients. All selected FASTQ files then underwent the same RNA-seq processing and analysis as FASTQ files generated from cell culture. Covariate data was also downloaded from the AMP-AD Knowledge Portal, and biological and technical covariates — sex, brain bank (referred to as “Source”), and the flowcell used for sequencing — were accounted for when fitting the model used by DESeq2. Examples of commands used to analyze all RNA-seq data are given in an additional file, along with quality control metrics that were organized using MultiQC 1.8 [5] (see S4 Code and Quality Control Data).

## Results

### Lentivirus infection of SH-SY5Y neuroblastoma cells yields isoform-specific overexpression of tau

To generate overexpression of non-mutant tau isoforms, we infected SH-SY5Y cells at an MOI of 20 with lentiviruses encoding 0N3R and 1N4R tau isoforms and differentiated cells with 15nM RA for 9 days. On the 8th day of differentiation, cells were treated either with 50 nM Aβ-42 oligomers to form a model of Alzheimer’s disease (characterized by overexpression of tau isoforms and presence of toxic Aβ species) or vehicle control (DMSO). For an overview of cell culture procedures, see Fig 1B.

After collection, cells underwent subcellular fractionation to observe tau expression in both the nucleus and cytoplasm, and Western blots were carried out using 3R and 4R tau isoform-specific antibodies to validate the experimental model (N=1). As expected, cell cultures treated with LV-3Rtau displayed overexpression of 3R tau relative to other groups, while those treated with LV-4Rtau showed distinct overexpression of 4R tau. Control groups displayed neither overexpression of 3R tau nor 4R tau. Histone 3 (H3) and GAPDH were used as loading controls for nuclear and cytoplasmic fractions, respectively (Fig 1A).

### Tau isoform overexpression and treatment with Aβ do not result in different nuclear/cytoplasmic total tau ratios

Loss of tau in the nucleus is associated with increased DNA damage and disruptions in heterochromatin organization [18, 37]. Moreover, there is evidence that Aβ may influence the localization of tau within neurons [40]. To ascertain whether a gain or loss of overall tau protein within the nucleus of differentiated SH-SY5Y cells occurred in the context of 3R or 4R tau overexpression and Aβ treatment, we performed immunofluorescence staining for total tau to assess potential different tau isoform-induced nuclear changes. Images were obtained with a confocal microscope and analysis was carried out using Fiji and R. To assess the translocation of tau into or out of the nucleus, we quantified the ratio of CTCF tau signal in the nucleus and in the cytoplasm. Overall, we found no significant differences in the ratio of tau expression in the nucleus or cytoplasm in tau-treated samples versus control samples (N=3). An additional file shows the results of this analysis (see S2 Fig).

### Tau isoform overexpression results in significantly fewer DNA double-strand breaks

Despite a lack of tau redistribution within differentiated SH-SY5Y cells, we found significant decreases in DNA double-strand breaks (DSBs) in tau-treated samples versus control samples (N=3) (Fig 2B). Notably, this phenomenon was most prevalent in samples treated with LV-4Rtau alone (p=0.037) or with Aβ (p=0.014) and completely absent in samples treated only with LV-3Rtau or with LV-3Rtau together with LV-4Rtau. Treatment with Aβ also appeared to have a slightly protective effect when given either to control samples (p=0.049) or to LV-3Rtau samples (p=0.014).

**Fig 2.**
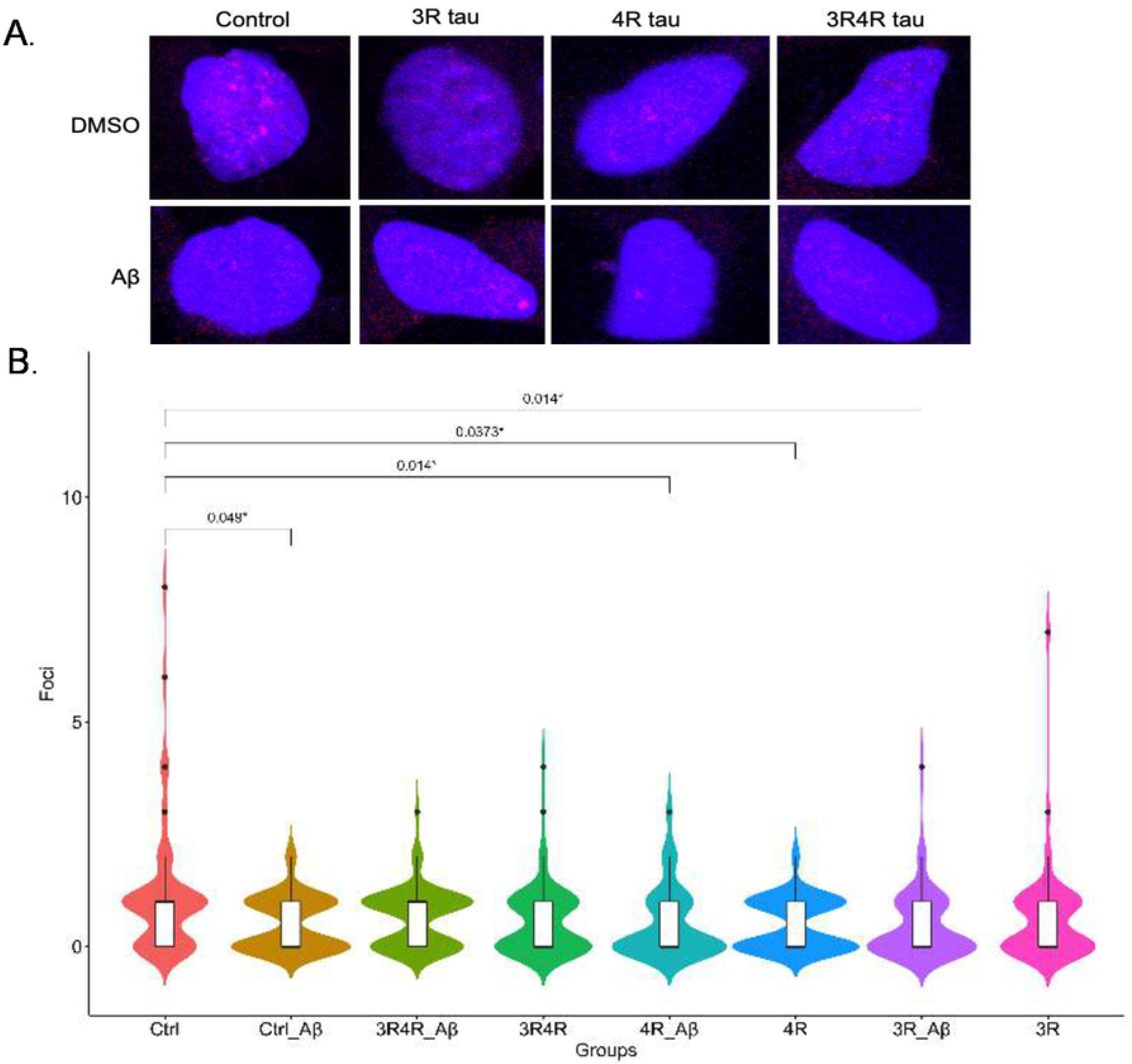
Overexpression of certain tau isoforms appears to lower DSB occurrence. (A) Confocal images (63x objective) of representative yH2AX (red) samples with nuclei stained with DAPI (blue). Red nuclear foci indicate where DSBs formed. (B) 4R tau overexpression, 3R tau overexpression with Aβ treatment and Aβ treatment of LV-Ctrl groups appear to correlate with smaller DSB distributions compared to control (N=3). All p values were calculated using a Wilcoxon ranked-sum test with multiple corrections done using FDR.

### Overexpression of 3R tau results in a consistent pattern of cell death

Next, we examined whether overexpression of tau isoforms affected cell survival. Using the CellTox Green Cytotoxicity Assay from Promega, which utilizes a green fluorescent dye to assess membrane damage (and thus cell nonviability), we quantified cell death across each experimental condition on each day of a typical differentiation procedure, excluding the first day when cells were infected and plated in 96-well plates. Because the fluorescent dye lasts for 72 hours, cells were segmented into three separate plates signifying three groups: Plate 1 measured cell death from the time lentiviruses were taken off the cells (Day 2) through Day 4 of the differentiation procedure, Plate 2 measured cell death from Days 5–7, and Plate 3 measured cell death as a result of Aβ treatment on each lentivirus condition (Days 8–9) (see Fig 3A for a schema of the workflow). Statistical analysis revealed a relatively consistent pattern of significant cell death associated with LV-3Rtau infection on most (6 out of 8) days measured (N=14); LV-3Rtau infection also showed higher levels of cell death compared to LV-4Rtau infection and 1:1 LV-3Rtau:LV-4Rtau infection groups, especially during Days 5–7 (Fig 3B). Notably, LV-4Rtau infections resulted in increases in cell death on Days 5, 6, and 9, while 1:1 LV-3Rtau:LV-4Rtau infections only showed increased cell death on Day 5. Aβ treatment did not appear to significantly affect cell viability in any group (Fig 3C–D). Although Aβ treatment appears to result in a significant increase in cell death in LV-3Rtau infections, this must be interpreted with caution because the vehicle (DMSO)-treated LV-3Rtau infection group shows significantly lower cell death from Days 8–9, implying that either Aβ does in fact have a deleterious interaction with 3R tau or that this result is an artefact of variation in the DMSO-treated group.

**Fig 3.**
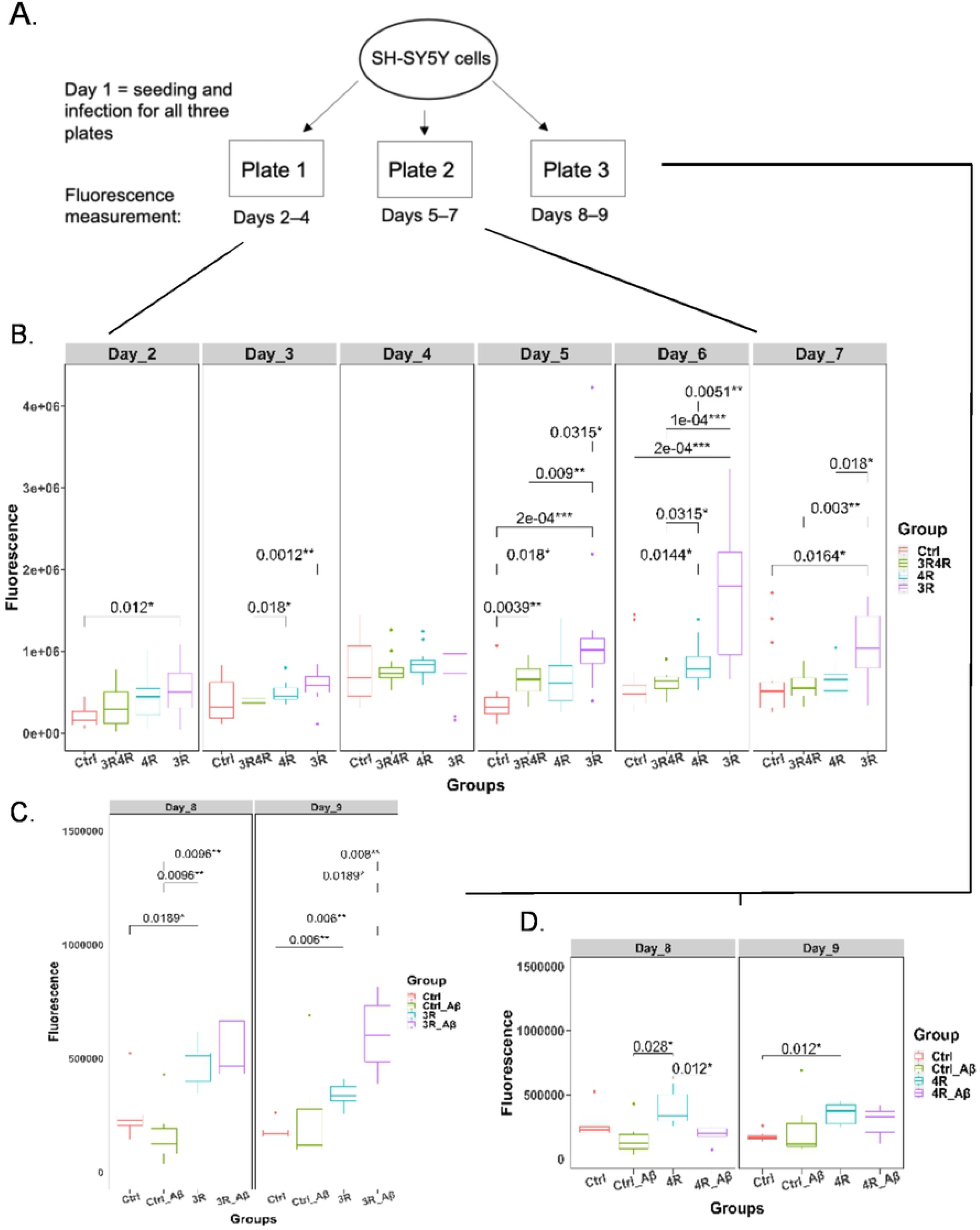
Cytotoxicity of tau isoforms and their interactions with Aβ. (A) Diagram mapping cytotoxicity assay workflow; plates 1 and 2 measured cytotoxicity through days 2–7, while plate 3 measured cytotoxicity on days 8–9, which captured cytotoxicity changes as a function of Aβ treatment. (B) Comparison of fluorescence values (cytotoxicity proxy, with higher fluorescence signifying greater levels of cell death) over days 2–7 for LV-Ctrl, LV-3R4Rtau, LV-4Rtau, and LV-3Rtau groups. (C) Comparison of fluorescence/cytotoxicity between LV-Ctrl treated with DMSO, LV-Ctrl treated with Aβ, LV-3Rtau treated with Aβ and LV-3Rtau treated with DMSO on days 8– 9, plate 3. (D) Comparison of fluorescence/cytotoxicity between LV-Ctrl treated with DMSO, LV-Ctrl treated with Aβ, LV-4Rtau treated with Aβ, and LV-4Rtau treated with DMSO on days 8–9, plate 3. Multiple comparison correction was done using FDR. Note: No significant changes were seen between LV-Ctrl treated with DMSO, LV-Ctrl treated with Aβ, LV-3R4Rtau treated with Aβ, and LV-3R4Rtau treated with DMSO on days 8– 9, plate 3. Data not shown. All p values were calculated using a Wilcoxon ranked-sum test with multiple corrections done using FDR.

### Tau isoforms, AD, and PSP produce uniquely altered gene expression

To examine potential mechanisms behind tau-mediated cellular dysfunction, we carried out 100 bp paired-end RNA-sequencing of cell culture samples (N=3). In addition, we analyzed 100 bp paired-end RNA-sequencing data from human patients with AD (N=60), PSP (N=71), and without any neurodegeneration (N=33) doi:10.1038/sdata.2016.89 [2]. Principal component analysis revealed distinct groupings between samples infected with tau lentiviruses and control samples, with less clear delineation between Aβ and vehicle (DMSO)-treated cultures (Fig 4A). AD patients’ data were similarly separated from control patients’ data, though PSP cases appeared less divided from controls (Fig 4B).

**Fig 4.**
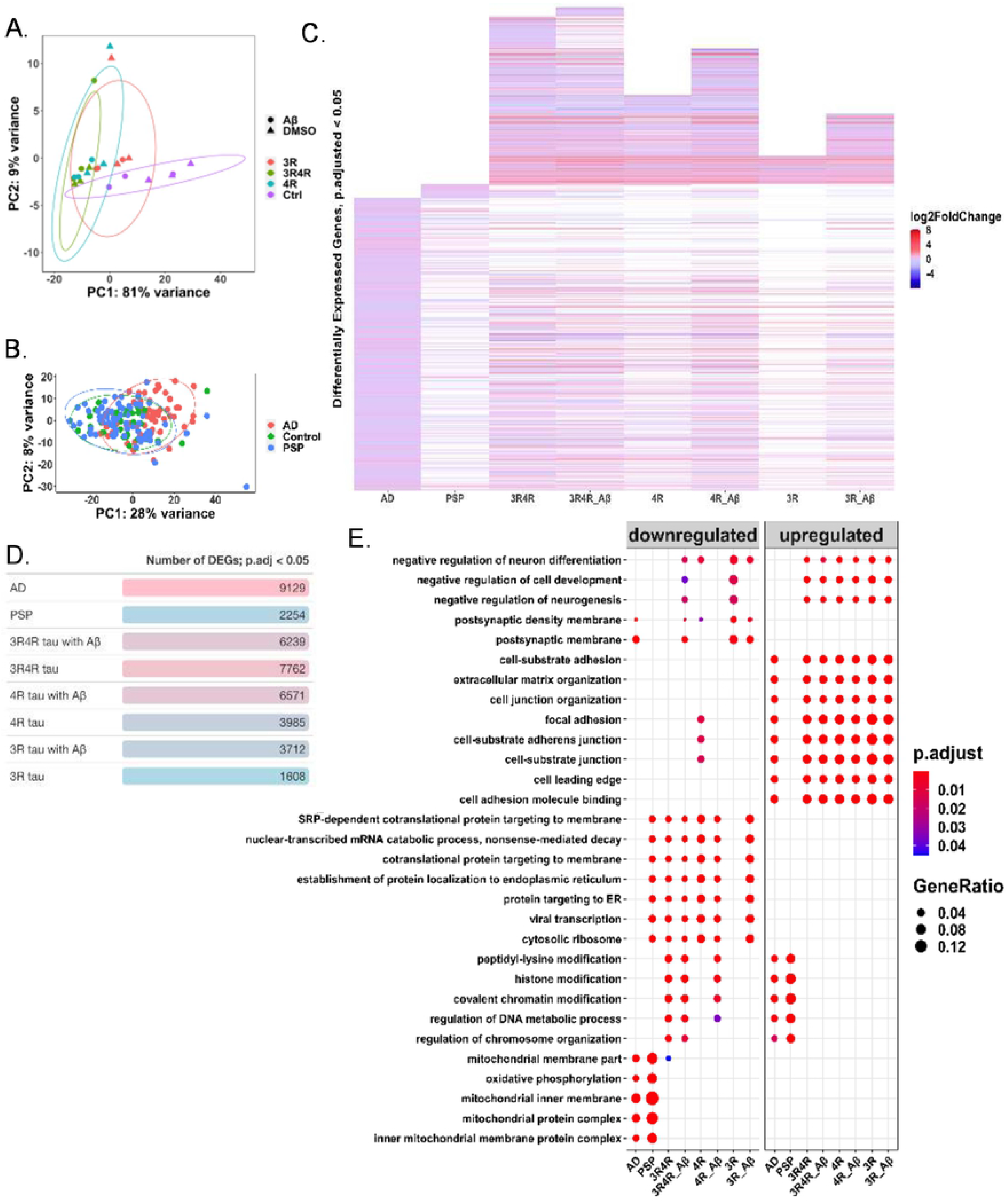
Patterns in differentially expressed genes in AD, PSP, and cell models. (A) PCA displaying grouping by gene expression between lentivirus and treatment conditions; PC1 accounts for 81% of variance between conditions and there is little separation between Aβ and DMSO-treated groups (N=3). (B) PCA displaying grouping by gene expression between AD (N=60), control (N=33), and PSP (N=71) patients. PC1 accounts for 28% of variation, with AD patients showing greater separation from control patients than PSP patients do. (C) Heatmap showing log-fold change in expression in genes considered significantly differentially expressed (FDR adjusted p-value<0.05) in each measured group (AD, N=60; PSP, N=71; cell groups, N=3 each) relative to their respective controls (control patients, N=33 for human disease cases; LV-Ctrl cells for cell conditions, N=3). White spaces indicate missing gene data, i.e., the genes were not significantly differentially expressed in these groups. LV-Ctrl treated with Aβ cell model is not shown due to too few significantly differentially expressed genes. (D) Table showing the number of significantly differentially expressed genes (FDR adjusted p-value<0.05) in each condition. (E) Overrepresentation analysis of gene-ontology enriched biological processes, molecular functions, and cellular components, with top 30 terms shown.

AD cases showed the greatest number of significantly differentially expressed genes, with PSP cases showing far fewer (Fig 4D). Among the cell culture samples, those infected with 1:1 LV-3Rtau:LV-4Rtau displayed the greatest number of differentially expressed genes (DEGs), followed by LV-4Rtau samples, and then LV-3Rtau samples. The samples infected with LV-Ctrl and treated with Aβ (the Aβ-only samples) showed the fewest significantly DEGs by far, with only 5 reaching the significance level of an adjusted p-value < 0.05. Overall, AD cases appeared to follow a distinct pattern of gene expression compared to the rest of the samples, while the cell culture samples significantly expressed a number of genes that did not meet the significance threshold in either PSP or AD (Fig 4C).

Gene expression patterns related to synaptic and mitochondrial functions were downregulated in AD, which concurs with studies that show dysfunction in these areas in AD patients [6, 21]. Upregulation of pathways pertaining to extracellular structure and focal adhesions were prevalent in all groups except PSP patients. Pathways related to protein synthesis appeared downregulated under all conditions except for AD patients and vehicle (DMSO)-treated LV-3Rtau samples. Downregulation of genetic and epigenetic phenomena occurred in both LV-3R:4Rtau samples and in LV-4Rtau samples, while these processes were upregulated in both AD and PSP. Finally, all cell culture groups displayed upregulation — with some also displaying downregulation — of processes related to cell development, negative regulation of neurogenesis, and negative regulation of neuron differentiation (Fig 4E).

### Tau isoforms, AD, and PSP produce distinct activation and repression of transposable elements

Recent studies suggest that tau may influence activation of TEs, which are mobile DNA sequences that have long been associated with genomic instability and more recently with potential regulatory functions [8, 12, 33]. To examine how tau isoforms might affect TE activation, we analyzed our RNA-seq data using software tools for identifying differentially expressed TE loci. PCA results show some separation between AD, PSP, and control patients (Fig 5B) and distinct groupings among cell culture conditions with small distances between DMSO and Aβ-treated samples (no TE loci were differentially expressed between any vehicle and Aβ-treated samples infected with the same lentivirus) (Fig 5A). Our analysis further revealed that overexpression of either tau isoform was sufficient to cause at least some TE dysregulation, though TEs appeared more abundantly dysregulated when 4R tau was overexpressed or when 3R tau was overexpressed in the context of Aβ treatment (Fig 5C).

**Fig 5.**
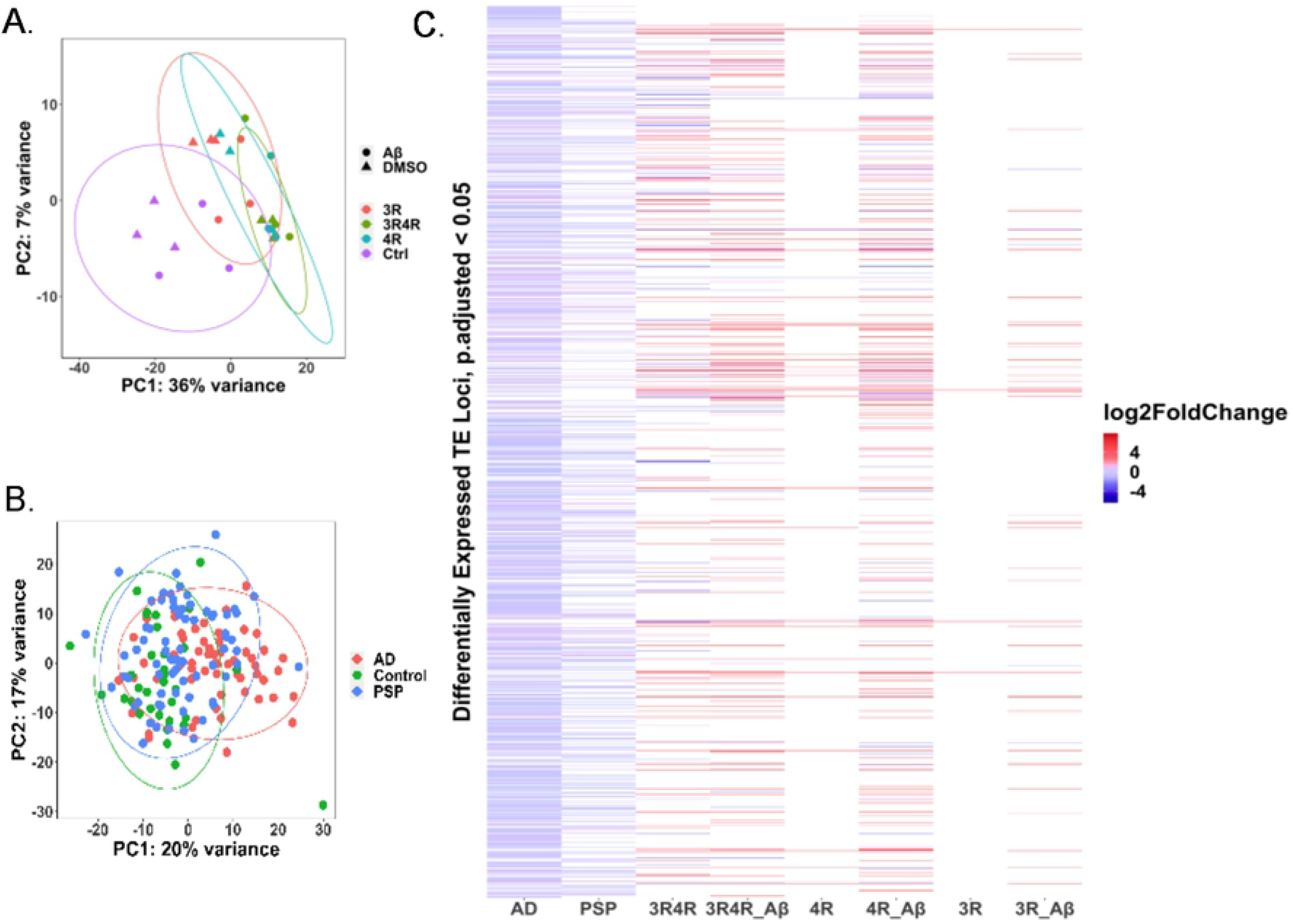
Patterns of locus-specific transposable element expression across human disease and cell samples. (A) PCA displaying grouping by locus-specific TE expression between lentivirus and treatment conditions; PC1 accounts for 36% of variation between conditions (N=3). (B) PCA displaying grouping by locus-specific TE expression between AD (N=60), control (N=33), and PSP (N=71) patients. PC1 accounts for 20% of variation, while PC2 accounts for 17% of variation. (C) Heatmap showing log-fold change in expression of TE loci considered significantly differentially expressed (FDR adjusted p-value<0.05) in each measured group (AD, N=60; PSP, N=71; cell groups, N=3 each) relative to their respective controls (control patients, N=33 for human disease cases; LV-Ctrl cells for cell conditions, N=3). White spaces indicate missing data, i.e., the TE loci were not significantly differentially expressed in these groups. LV-Ctrl treated with Aβ cell model did not display any differentially expressed TE loci relative to LV-Ctrl and is therefore not included in the heatmap.

The two families of TEs that represent the most dysregulated loci were the L1 and Alu families, which are classified as part of the LINE and SINE superfamilies, respectively (Fig 6A–B). The L1 family in particular is considered the only autonomous TE family still active in the human genome, while Alu can be activated by L1 activity [35]. Far greater numbers of TEs showed differential expression in AD and PSP cases compared to cell culture samples, likely reflecting the greater cellular heterogeneity and complexity of bulk RNA-seq data from human brains than from cell culture (Fig 5C, 6A– B). AD and PSP displayed similar patterns of TE expression, though more TE loci were differentially expressed in AD than in PSP (Fig 5C, 6A–B). No TEs were differentially expressed in Aβ-treated LV-Ctrl cell cultures, suggesting that TE activation is not solely driven by Aβ, but may be promoted by tau pathology, confirming the results of other studies [8]. TE activation and repression also appear to show patterns of expression throughout the autosomal and X chromosomes depending on which tau isoform is overexpressed and whether this overexpression is accompanied by treatment with Aβ (Fig 6C). Finally, AD and PSP show the greatest number of differentially expressed TE families, with AD showing the highest amount. Among the cell culture groups, LV-3R4R samples showed the most differentially expressed TE families. LV-3R samples treated with DMSO showed no overall differentially expressed TE families (Fig 6D).

**Fig 6.**
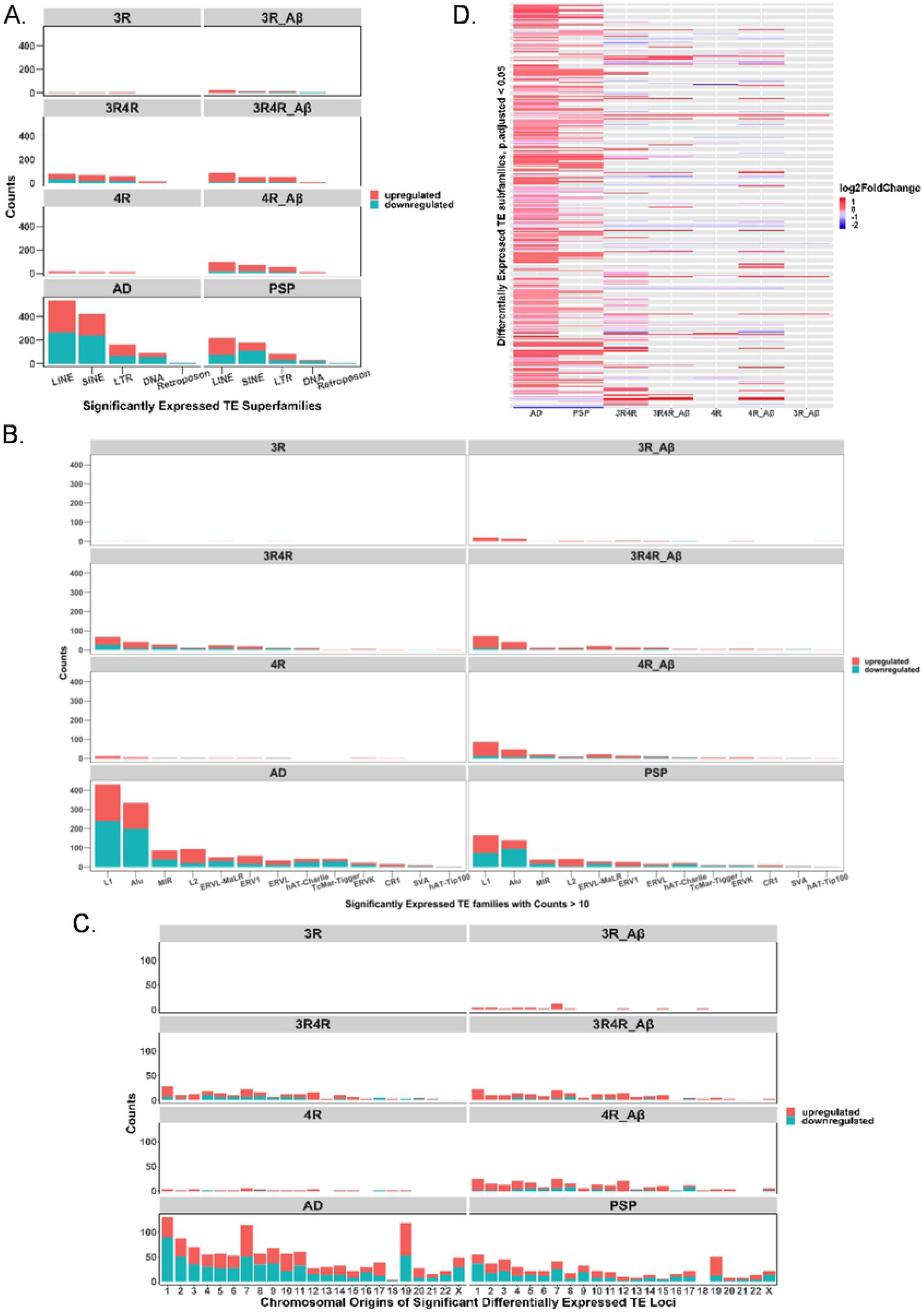
Classification of significantly differentially expressed locus-specific transposable elements by transposable element superfamily and family. (A) Counts of significantly differentially expressed TE loci (downregulated and upregulated) per condition grouped by superfamily of TE. (B) Counts (of families with counts > 10) of significantly differentially expressed TE loci (downregulated and upregulated) per condition grouped by TE family. (C) Chromosomal distribution of upregulated and downregulated TE loci on autosomal and X chromosomes. (D) Heatmap of differentially expressed TE subfamilies (i.e., viewing TE expression at subfamily level instead of locus-level; see supporting information for lists of specific subfamilies in AD and PSP (S3 Table). Note that this heatmap was generated from counts calculated using TEcounts, which by default provides data on subfamilies, and not TElocal).

## Discussion

Despite being one of the leading causes of death worldwide, neurodegenerative diseases are notoriously lacking in any kind of therapeutic treatment. Though several toxic forms of proteins are major actors in these types of diseases, abnormal expression of one or more tau protein isoforms is characteristic of a significant portion of them. Mechanisms underlying tau-mediated neurodegeneration and the unique roles of its isoforms remain poorly understood, and recent studies have shown that tau may play more diffuse roles in the cell than binding to microtubules. Creating a fuller picture of how tau and its isoforms may lead to neurodegeneration is thus critical for designing treatments for tau-based diseases.

In this study, we sought to elucidate differences in how two major types of isoforms, 3R and 4R tau, promote pathways to cellular dysfunction. By analyzing the transcriptomes in a cell culture model of tau isoform overexpression, we demonstrated that 3R and 4R tau isoforms, when overexpressed alone or in combination with each other and with Aβ, promoted markedly different transcriptomic patterns. Notably, while no combination of tau isoform overexpression and Aβ is sufficient to recapitulate all dysregulated pathways seen in our analysis of RNA-seq data from AD and PSP patients, overexpression of either isoform is sufficient to cause dysregulation in TE expression, though this effect is more prominent with 4R tau overexpression. Only when Aβ was introduced did the 3R tau samples start to replicate the patterns of TE expression seen in AD, PSP, and the other cell samples. Interestingly, the LV-Ctrl group treated with Aβ alone failed to show any dysregulated TE expression, implying that while Aβ alone may not be sufficient to drive TE dysregulation, it appears to aggravate aberrant TE expression in the context of tauopathy. This pattern of dysregulated TE expression in cells also paralleled our findings in the RNA-seq analyses in the clinical samples in which both AD and PSP patients samples had dysregulated TE expression but the AD patient samples appeared to have higher dysregulated TE expression than the PSP patient samples. This is consistent with greater dysregulation of TE expression with 3R4R tau in the presence of Aβ as featured in AD but to a lesser degree with PSP (4R tau in the absence of Aβ). Recent evidence shows that tau pathology can broadly impact the epigenome through heterochromatin relaxation and histone acetylation [7, 8, 12]. Mechanistically, then, it stands to reason that tau-driven epigenetic changes could lead to TE dysregulation [8, 33]. To our knowledge, our study is the only one to examine the transcriptomes of overexpressed 3R and 4R tau isoforms in interaction with Aβ, and to show a tau isoform-dependent change in TE dysregulation. TE dysregulation has only recently been discovered in tauopathies, and has been garnering attention in other neurodegenerative diseases [34].

Despite the current lack of research on TEs in neurodegeneration, one study has shown that in a tauopathy model of Drosophila, treatment with a reverse transcriptase inhibitor reduces TE activation and increases longevity [33]. In our study, the two TE families with the greatest number of dysregulated loci were the L1 and Alu families. Intriguingly, these two families are main suspects in the pathogenesis of other neurological diseases. Both, for instance, are implicated in Aicardi-Goutieres Syndrome, which is characterized by neuroinflammation driven by increased type I-interferon activity [34]. Links between L1 expression and the dedifferentiation of cells in AD have also been drawn [34]. If TE dysregulation does in fact promote neurological damage in a variety of diseases, this could provide a novel and actionable avenue for therapeutic interventions.

Aside from dysregulation in TE expression, several notable phenomena emerged from our RNA-seq analysis. Gene Ontology (GO) overrepresentation analysis revealed downregulation of several processes related to protein synthesis and the ribosome in PSP and most other cell samples. Tau-mediated disruption in protein synthesis has been previously noted to be a contributing factor to neurodegeneration [19]. Terms related to structural components of the cell, such as the extracellular matrix and adherence junctions, were also largely upregulated in AD and the cell groups, as might be expected given the role of tau in cytoskeletal organization. As in TE expression, the presence of Aβ appears to drive the LV-3Rtau group toward transcriptomic patterns that converge with those of LV-4Rtau and LV-3R4Rtau groups.

Other aspects of our study also showed interesting divergence in tau isoform-driven effects. For instance, 3R tau consistently showed higher cytotoxicity levels compared to control groups than did other samples, and occasionally showed higher toxicity than either the LV-3R4R or LV-4R groups. These results concur with literature suggesting that the 0N3R tau isoform led to shorter lifespans in transgenic Drosophila [27], but contrast with other studies which conclude that 4R tau overexpression is more pathogenic, especially in htau transgenic mice [26]. While greater clarification is needed on this issue, the existence of both 3R and 4R tauopathies suggest that overexpression of either isoform is inherently toxic. Ultimately, researching mechanisms of how these isoforms produce cytotoxicity may prove more fruitful, and recent research has illuminated how overexpression of either isoform can lead to opposite disruptions in vesicle transport [14]. More surprisingly, the 1:1 combination of 3R and 4R tau in our model showed relatively low levels of cytotoxicity compared to controls. Relatively high spread in the distributions of all groups may partially account for this, and either a more sensitive assay or higher numbers of samples may be called for when assessing cytotoxicity in response to these isoforms over several days. Alternatively, it may be that the equal overexpression of both isoforms requires more time to show its cytotoxic effects; the LV-3R4R tau groups showed much greater transcriptional dysregulation than the other groups, which suggests that these groups are in fact being affected by their overexpression of both isoforms. Notably, these samples also showed the strongest dysregulation of TEs among all cell culture groups and followed a similar pattern of TE disruption found in AD and PSP. Whether these transcriptional changes eventually manifest in greater cytotoxicity remain to be seen.

We also sought to determine whether isoform-specific tau overexpression had any effect on the prevalence of DSBs, which are highly damaging to cells and have been found to be increased in the brains of people with AD and mild cognitive impairment [28]. Tau deficiency has been previously linked to DNA damage, and overexpression of tau in tau-deficient models has been shown to decrease DNA damage [32, 37]. Though our data was characterized by high levels of variation and outliers, we found that overexpression of the 4R tau isoform in particular resulted in fewer DSBs as measured by yH2AX fluorescence. Notably, changes in DSBs in our cell models could not be correlated with any change in the ratio of nuclear vs. cytoplasmic levels of tau, as no samples showed any significant translocation of tau, although we did not assess whether there was any change in localization of tau isoforms. This result did not exclude the possibility of tau isoforms affecting DNA integrity or chromatin organization, as more subtle pathways affecting these components may be involved, but led us to conclude that relative tau distribution from the nucleus to the cytoplasm is unaffected by Aβ treatment or overexpression of certain tau isoforms.

Overall, our study illuminates several distinctions in how 3R and 4R tau isoforms may disrupt normal cellular function and reveals that TE dysregulation can result from overexpression of non-mutant human tau isoforms, with expression patterns recapitulating those of AD and PSP. Our study shows both locus-specific TE and global TE expression patterns in context of tau isoform overexpression and in AD and PSP cases. The only other two studies to examine TE expression in AD have concluded that LTR families of TEs are the most dysregulated; however, these studies depended on taking a global view of TE expression, instead of a locus-specific one (it should be noted that software for locus-specific TE detection was likely not available for these studies, as it has only recently been developed). Our own global view of TE expression largely confirms the pattern of their results, yet interestingly, when viewed from a locus-specific standpoint, L1 and Alu families are far better represented among the dysregulated TEs. This is particularly notable given that L1 in particular is the only active and autonomous TE family in humans, while Alu is an active element that hijacks L1 replicative machinery. These results will eventually need more stringent validation than is currently available; repetitive DNA regions, which describe most TEs, are notoriously difficult to call in current short-read sequencing platforms, and long-read platforms, while promising, have issues with error rates and read depth to overcome [23]. Since both the number of dysregulated TE loci and the number of dysregulated TE families remains much higher in AD and PSP than in our cell culture models, it is likely that there are other mechanisms also at play in TE expression in disease. Because our cell model only evaluated neuronal cells, it would be illuminating to see how TE expression may differ among the other cell types known to be affected by aberrant tau. Moreover, the link between tau pathology and epigenetic changes provides a possible mechanism for tau-induced TE differential expression, meriting a more data-intensive look at how tau and its isoforms affect chromatin remodeling than has previously been performed. Overall, the evidence supporting the notion that TE dysregulation could be a cytotoxic, therapeutically targetable consequence of pathogenic tau offers a new, exciting vantage point into the nature of tauopathic diseases.

## Conclusions

In this study, we provide insight into how 3R and 4R tau isoforms uniquely affect the transcriptome and cell death. We show that overexpression of these isoforms, especially 4R tau, are sufficient to produce differential expression of transposable elements, a therapeutically targetable, proposed source of inflammation and cause of cell death which has only recently been discovered in the context of tauopathies. Furthermore, we used newly available software to map the differentially expressed transposable elements in both our cell lines and in human tissue to specific locations within the genome, a level of detail that has not been previously reported on. In doing so, we show that while family-level counts of transposable element expression show more significant differences in LTRs and hERVs, LINEs and SINEs, which represent relatively autonomous transposable elements that have been previously associated with other neurological disorders, appear more widely dysregulated at the locus-level. Though current technology limits how confident we can be in determining expression levels of repeat elements, these results bolster support for future investigation into the role of transposable element expression in neurodegenerative diseases.

## Acknowledgements

The authors thank the members of the Rissman lab at UCSD for technical help and support.

## Supporting information

**S1_raw_images**. Original blots used for Fig. 1A.

**S2 Fig. Total tau ratio between nucleus and cytoplasm is unchanged by overexpression of tau isoforms and Aβ**. Immunofluorescence data showing ratio of total tau expression in the nucleus or cytoplasm in tau-treated samples versus control samples (N=3).

**S3 Table. Significantly differentially expressed genes and transposable elements from AD and PSP patient data**. Excel document listing multiple tables with differentially expressed genes (determined through featureCounts and DESeq2) and transposable element loci and families found using TElocal and TEcount, respectively. Significance is defined by FDR-adjusted p values < 0.05.

**S4 Code and Quality Control Data**. ZIP file containing examples of code used in analyzing RNA-seq data along with MultiQC reports of RNA-seq quality control data from cells.

## References

1. Adams SJ, DeTure MA, McBride M, Dickson DW, Petrucelli L. Three repeat isoforms of tau inhibit assembly of four repeat tau filaments. PLoS One. 2010 May 25;5(5):e10810.

2. Allen M, Carrasquillo MM, Funk C, Heavner BD, Zou F, Younkin CS, et al. Human whole genome genotype and transcriptome data for Alzheimer’s and other neurodegenerative diseases. Sci Data. 2016 Oct 11;3:160089.

3. Andrews S. FastQC: a quality control tool for high throughput sequence data. 2010.

4. Dobin A, Davis CA, Schlesinger F, Drenkow J, Zaleski C, Jha S, et al. STAR: ultrafast universal RNA-seq aligner. Bioinformatics. 2013 Jan 1;29(1):15–21.

5. Ewels P, Magnusson M, Lundin S, Käller M. MultiQC: summarize analysis results for multiple tools and samples in a single report. Bioinformatics. 2016 10 1;32(19):3047–8.

6. Forner S, Baglietto-Vargas D, Martini AC, Trujillo-Estrada L, LaFerla FM. Synaptic Impairment in Alzheimer’s Disease: A Dysregulated Symphony. Trends Neurosci. 2017 06;40(6):347–57.

7. Frost B, Hemberg M, Lewis J, Feany MB. Tau promotes neurodegeneration through global chromatin relaxation. Nat Neurosci. 2014 Mar;17(3):357–66.

8. Guo C, Jeong HH, Hsieh YC, Klein HU, Bennett DA, De Jager PL, et al. Tau Activates Transposable Elements in Alzheimer’s Disease. Cell Rep. 2018 06 5;23(10):2874–80.

9. Guo T, Noble W, Hanger DP. Roles of tau protein in health and disease. Acta Neuropathol.2017 05;133(5):665–704.

10. Jin Y, Hammell M. Analysis of RNA-Seq Data Using TEtranscripts. Methods Mol Biol. 2018;1751:153–67.

11. Jin Y, Tam OH, Paniagua E, Hammell M. TEtranscripts: a package for including transposable elements in differential expression analysis of RNA-seq datasets. Bioinformatics. 2015 Nov 15;31(22):3593–9.

12. Klein HU, McCabe C, Gjoneska E, Sullivan SE, Kaskow BJ, Tang A, et al. Epigenome-wide study uncovers large-scale changes in histone acetylation driven by tau pathology in aging and Alzheimer’s human brains. Nat Neurosci. 2019 01;22(1):37–46.

13. Klein SJ, O’Neill RJ. Transposable elements: genome innovation, chromosome diversity, and centromere conflict. Chromosome Res. 2018 03;26(1-2):5–23.

14. Lacovich V, Espindola SL, Alloatti M, Pozo Devoto V, Cromberg LE, Carná ME, et al. Tau Isoforms Imbalance Impairs the Axonal Transport of the Amyloid Precursor Protein in Human Neurons. J Neurosci. 2017 01 4;37(1):58–69.

15. Lebouvier T, Pasquier F, Buée L. Update on tauopathies. Curr Opin Neurol. 2017 Dec;30(6):589–98.

16. Liao Y, Smyth GK, Shi W. featureCounts: an efficient general purpose program for assigning sequence reads to genomic features. Bioinformatics. 2014 Apr 1;30(7):923–30.

17. Love MI, Huber W, Anders S. Moderated estimation of fold change and dispersion for RNA-seq data with DESeq2. Genome Biol. 2014;15(12):550.

18. Mansuroglu Z, Benhelli-Mokrani H, Marcato V, Sultan A, Violet M, Chauderlier A, et al. Loss of Tau protein affects the structure, transcription and repair of neuronal pericentromeric heterochromatin. Sci Rep. 2016 09 8;6:33047.

19. Meier S, Bell M, Lyons DN, Rodriguez-Rivera J, Ingram A, Fontaine SN, et al. Pathological Tau Promotes Neuronal Damage by Impairing Ribosomal Function and Decreasing Protein Synthesis. J Neurosci. 2016 Jan 20;36(3):1001–7.

20. Muñoz-López M, García-Pérez JL. DNA transposons: nature and applications in genomics. Curr Genomics. 2010 Apr;11(2):115–28.

21. Moreira PI, Carvalho C, Zhu X, Smith MA, Perry G. Mitochondrial dysfunction is a trigger of Alzheimer’s disease pathophysiology. Biochim Biophys Acta. 2010 Jan;1802(1):2–10.

22. Okonechnikov K, Conesa A, García-Alcalde F. Qualimap 2: advanced multi-sample quality control for high-throughput sequencing data. Bioinformatics. 2016 Jan 15;32(2):292–4.

23. O’Neill K, Brocks D, Hammell MG. Mobile genomics: tools and techniques for tackling transposons. Philos Trans R Soc Lond B Biol Sci. 2020 03 30;375(1795):20190345.

24. Payer LM, Burns KH. Transposable elements in human genetic disease. Nat Rev Genet. 2019 12;20(12):760–72.

25. Rockenstein E, Overk CR, Ubhi K, Mante M, Patrick C, Adame A, et al. A novel triple repeat mutant tau transgenic model that mimics aspects of pick’s disease and fronto-temporal tauopathies. PLoS One. 2015;10(3):e0121570.

26. Schoch KM, DeVos SL, Miller RL, Chun SJ, Norrbom M, Wozniak DF, et al. Increased 4R-Tau Induces Pathological Changes in a Human-Tau Mouse Model. Neuron. 2016 06 1;90(5):941–7.

27. Sealey MA, Vourkou E, Cowan CM, Bossing T, Quraishe S, Grammenoudi S, et al. Distinct phenotypes of three-repeat and four-repeat human tau in a transgenic model of tauopathy. Neurobiol Dis. 2017 Sep;105:74–83.

28. Shanbhag NM, Evans MD, Mao W, Nana AL, Seeley WW, Adame A, et al. Early neuronal accumulation of DNA double strand breaks in Alzheimer’s disease. Acta Neuropathol Commun. 2019 05 17;7(1):77.

29. Sotiropoulos I, Galas MC, Silva JM, Skoulakis E, Wegmann S, Maina MB, et al. Atypical, non-standard functions of the microtubule associated Tau protein. Acta Neuropathol Commun. 2017 Nov 29;5(1):91.

30. Spencer B, Michael S, Shen J, Kosberg K, Rockenstein E, Patrick C, et al. Lentivirus mediated delivery of neurosin promotes clearance of wild-type α-synuclein and reduces the pathology in an α-synuclein model of LBD. Mol Ther. 2013 Jan;21(1):31–41.

31. Stine WB, Jungbauer L, Yu C, LaDu MJ. Preparing synthetic Aβ in different aggregation states. Methods Mol Biol. 2011;670:13–32.

32. Sultan A, Nesslany F, Violet M, Bégard S, Loyens A, Talahari S, et al. Nuclear tau, a key player in neuronal DNA protection. J Biol Chem. 2011 Feb 11;286(6):4566–75.

33. Sun W, Samimi H, Gamez M, Zare H, Frost B. Pathogenic tau-induced piRNA depletion promotes neuronal death through transposable element dysregulation in neurodegenerative tauopathies. Nat Neurosci. 2018 08;21(8):1038–48.

34. Tam OH, Ostrow LW, Gale Hammell M. Diseases of the nERVous system: retrotransposon activity in neurodegenerative disease. Mob DNA. 2019;10:32.

35. Terry DM, Devine SE. Aberrantly High Levels of Somatic LINE-1 Expression and Retrotransposition in Human Neurological Disorders. Front Genet. 2019;10:1244.

36. Tiscornia G, Singer O, Verma IM. Production and purification of lentiviral vectors. Nat Protoc. 2006;1(1):241–5.

37. Violet M, Delattre L, Tardivel M, Sultan A, Chauderlier A, Caillierez R, et al. A major role for Tau in neuronal DNA and RNA protection in vivo under physiological and hyperthermic conditions. Front Cell Neurosci. 2014;8:84.

38. Xicoy H, Wieringa B, Martens GJ. The SH-SY5Y cell line in Parkinson’s disease research: a systematic review. Mol Neurodegener. 2017 01 24;12(1):10.

39. Yu G, Wang LG, Han Y, He QY. clusterProfiler: an R package for comparing biological themes among gene clusters. OMICS. 2012 May;16(5):284–7.

40. Zempel H, Thies E, Mandelkow E, Mandelkow EM. Abeta oligomers cause localized Ca(2+) elevation, missorting of endogenous Tau into dendrites, Tau phosphorylation, and destruction of microtubules and spines. J Neurosci. 2010 Sep 8;30(36):11938–50.

